# Identification of Pesticides Associated with an Increased Risk of Parkinson’s Disease using a Multi-Screen Approach

**DOI:** 10.1101/2025.07.22.666186

**Authors:** Marisol Arellano, Lisa M Barnhill, Aaron M Kim, Kazi Md Mahmudul Hasan, Sharon Li, Kimberly C Paul, Chao Peng, Beate Ritz, Jeff M Bronstein

## Abstract

Parkinson’s Disease (PD) is a progressive neurodegenerative disease characterized by aggregation and transmission of alpha-synuclein (α-syn) protein and loss of dopaminergic neurons. The etiology of PD is multifactorial involving both genetic and environmental factors. Pesticide exposure has been associated with PD, but it is still unclear which of the hundreds of chemically and structurally diverse pesticides confer this association. While there are numerous pesticides currently registered for use in the United States, in this study, we focused on 62 pesticides used in the California Central Valley. We used 2 cell-based screens and a recently developed pesticide-wide association study (PWAS) analysis to identify candidates that increase the risk of PD. One cell-based screen tested for pesticides that alter autophagy and the other tested for their effect on α-syn transmission. To establish biological plausibility, pesticides that were positive in all 3 screens were tested for dopaminergic neurotoxicity in an *in vivo* zebrafish (ZF) model. The autophagy screen resulted in a total of 22 hits for alterations to either autophagosomes (16 hits) and lysosomes (14 hits) out of the 62 pesticides tested. The α-syn transmission screen resulted in 29 hits, and the PWAS resulted in 34 hits. Six pesticides were positive in all 3 screens and four of these pesticides induced aminergic neuron loss in ZF larvae. The majority of the pesticides identified in our screens have not previously been implicated as risk factors for PD but should be considered in future studies.

## 1. Introduction

Parkinson’s Disease (PD) is an increasingly prevalent neurodegenerative disorder globally, and the number of cases is expected to grow with the aging population^1,2^. The hallmarks of PD include pathological aggregation of α-synuclein (α-syn) protein in Lewy bodies, progressive loss of dopaminergic neurons in the substantia nigra, and neuroinflammation that lead to progressive motor and cognitive impairments^3–6^. The etiology of PD is complex and involves multiple factors, including genes and the environment. The earliest pathology of aggregated α- syn is found in the gut and olfactory bulb and is believed to spread trans-synaptically throughout the central nervous system (CNS).

α-syn plays an important role in the regulation of synaptic vesicle trafficking and fusion at the presynaptic terminal but some aggregated forms of α-syn are toxic to neurons. Furthermore, insoluble α-syn has been shown to have seeding capability with monomeric α-syn, which can result in the spread of pathological α-syn species^7^. Studies evaluating the transmissibility of pathogenic species of α-syn protein suggest that the intercellular spread of these pathological seeds precedes the templated amplification of misfolded α-syn in recipient cells to form misfolded protein inclusions that are resistant to enzymatic breakdown^8,9^.

It is not well understood what initiates α-syn aggregation, but the identification of rare genetic forms of PD have provided important clues into its pathophysiology. Increased expression of α-syn arising from gene multiplication is sufficient to cause PD^10–15^. Furthermore, polymorphisms in the α-syn gene (*SNCA*) are associated with increased risks of developing PD and faster symptom progression in those who develop PD^16–18^. Other gene mutations leading to PD have implicated disruption of proteostasis as an adverse outcome pathway in at least some familial cases^12,19–21^. For example, mutations in *GBA*, *LRRK2*, *VPS35*, *RAB32*, and *ATP13A2* lead to altered autophagy, a primary mechanism for clearing α-syn aggregates. Further support for altered autophagy as a pathogenic pathway to developing PD comes from environmental studies. Air pollution has recently been identified as a risk factor for PD and diesel exhaust extracts have been reported to produce PD pathology by disrupting autophagy^22–24^. Thus, increased levels of α- syn either through increased expression or decreased degradation can lead to the development of PD.

Since genetic factors alone account for a minority of PD cases, it is likely that environmental factors also contribute considerably to PD risk. As the environment is modifiable, it is imperative to identify specific environmental factors that alter the risk of disease and determine the mechanisms by which they act. Pesticide exposure is one of the most well-documented environmental risk factors associated with increasing risk of PD^25–27^. Pesticides are chemicals used to control, repel, or kill pests, such as insects, rodents, fungi, and weeds. They are widely used in the United States, both commercially and privately, and are composed of hundreds of different chemicals^27–29^. Several studies have reported the association between pesticides and PD, but only a few specific chemicals have been identified that confer increased risk. We recently assessed the relationship between individual pesticides and PD utilizing population-based case-control study data and an untargeted approach for pesticide exposures ^30^. To determine which of the specific pesticides identified epidemiologically are associated with PD, here we used 2 cell-based screens to identify pesticides that alter autophagy and α-syn transmission and an epidemiologic screen using a pesticide exposure-wide association study (PWAS) to prioritize pesticides for further investigation (Figure 1). The intent of this approach is to identify pesticides most likely to be altering PD risk with the understanding that it will not be all-inclusive since there are likely other pathological pathways involved in the pathogenesis of PD. Despite this limitation, we identified 6 specific pesticides that altered α-syn transmission and autophagy that were initially also associated with PD in the untargeted analysis. Additional biological plausibility was established in 4 of the 6 as they were toxic to dopaminergic neurons in a *Danio rerio* zebrafish (ZF) model.

**Figure 1.**
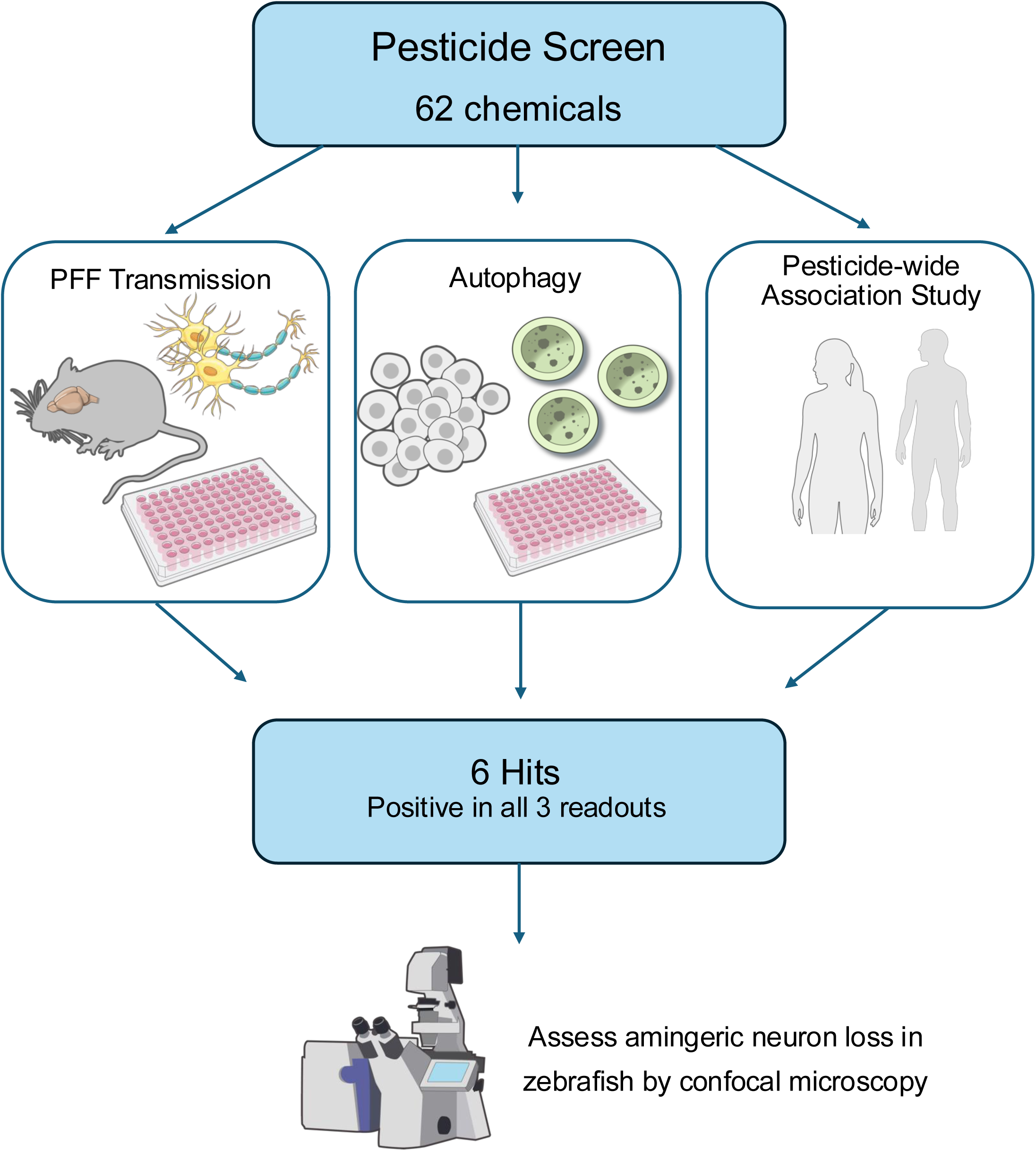
Pesticide Screen Schematic. An overview of the organization of the screen to evaluate candidate pesticides on key adverse outcome pathways involved in Parkinson’s Disease (PD) and on individual chemical association with PD using PEWAS. Pesticide candidates are funneled through each phase independently, and the results are collated to identify hits that are positive in all 3 readouts. The hits are then evaluated for the effect on aminergic neuron loss in a ZF model of neurodegeneration.

## 2. Methods

### 2.1 Chemicals and reagents

Chemicals were primarily purchased from ChemService (West Chester, PA, USA) and Sigma-Aldrich (St. Louis, MO, USA). A full list of the chemicals can be found in Supplement Table S1. Information for reagents purchased for experiments is listed below.

### 2.2 Pesticide selection

Pesticide selection for the screen was informed by reported agricultural use in California’s Central Valley and exposure to the patients and controls of Parkinson’s disease within the Parkinson, Environment, and Gene (PEG) study cohort. California law mandates the recording of all commercial agricultural pesticide applications, allowing us to track the individual pesticide active ingredients that have been applied near the homes and workplaces of the PEG study participants since 1974^30^. Overall, 722 different chemicals were applied within a 500m buffer of at least one PEG participant’s residence or workplace. We aimed to agnostically assess chemicals in relation to PD and therefore included as many pesticides as possible while balancing for the labor-intensive assays conducted in the study. The pesticide screen panel was limited to pesticides for which we found that greater than 10% of the participants had been exposed and among these 137 pesticides, we included at least 1 pesticide from each chemical class. In total, 62 pesticides met these criteria (Supplement Table S1) and were incorporated into the pesticide panel to be screened.

### 2.3 Animal husbandry and care

Embryos from gestating CD1 mice (Charles River Laboratories, Thousand Oaks, CA, USA) were used for primary neuron cultures; dams were maintained on a standard 12-hour light/ 12-hour dark cycle with *ad libitum* access to food and water.

ZF (*Danio rerio*) were raised at 28°C in recirculating water tanks on a 14-hour light / 10-hour dark cycle and were fed twice a day with brine shrimp. Embryos and larvae were obtained from natural mating and staged according to days post fertilization (dpf). ET*vmat2*:eGFP ZF, which expresses eGFP in aminergic neurons under the vesicular monoamine transporter (*vmat2)* promoter, were used for neuronal quantification.

All breeding, housing, and experimental procedures adhered to the NIH Guide for the Care and Use of Experimental Animals and were approved by the UCLA Institutional Animal Care and Use Committee (IACUC).

### 2.4 Fibril preparation

Purification of recombinant human α-syn and generation of α-syn PFFs were performed as previously described^31^. Briefly, the pRK172 plasmid containing the mouse *SNCA* gene was transformed into BL21 (DE3) competent E. *coli* (C2527H, New England). A single colony was expanded in Terrific Broth (12 g/L Bacto-tryptone, 24 g/L yeast extract, 4 mL/L glycerol, 9.4 g/L KH₂PO₄, and 2.2 g/L K₂HPO₄) supplemented with ampicillin.

Bacterial pellets were collected, sonicated, and boiled to precipitate unwanted proteins. The supernatant was dialyzed overnight in 10 mM Tris (pH 7.6), 50 mM NaCl, and 1 mM EDTA. The protein solution was then filtered through a 0.22 µm filter and concentrated using Pierce Protein Concentrators (PI88517, Thermo Scientific). The concentrated protein was loaded onto a Superdex 200 column (28990944, Cytiva), and 1 mL fractions were collected. These fractions were analyzed by SDS-PAGE and Coomassie blue staining, and those highly enriched in α-syn were pooled and dialyzed overnight in 10 mM Tris (pH 7.6), 25 mM NaCl, and 1 mM EDTA.

The dialyzed protein was subjected to anion exchange chromatography using a HiTrap Q HP column (17115401, Cytiva) with a linear gradient from 25 mM to 1 M NaCl. Collected fractions were analyzed by SDS-PAGE and Coomassie blue staining, and those enriched in α-syn were pooled, dialyzed into DPBS, filtered through a 0.22 µm filter, and concentrated to >5 mg/mL using Pierce Protein Concentrators. The purified monomer was aliquoted and stored at -80°C. To generate α-syn PFFs, 5 mg/mL α-syn monomer was incubated at 37°C with shaking at 1,000 rpm for seven days.

### 2.5 Primary neuron isolation

Mouse hippocampal neurons were prepared in-house from E16.5 embryos of CD1 mice (Charles River) as previously described^8,31,32^. Prior to cell plating, 384-well plates (781091, Greiner Bio-One) were coated with 0.1 mg/mL Poly-D-lysine (P0899, Sigma-Aldrich) in 50 mM borate buffer (pH 8.5) and incubated overnight at room temperature. The following day, plates were washed five times with ddH₂O before use.

On the day of dissection, E16.5 embryos were extracted from the uterus and decapitated. The heads were rinsed four times in ice-cold 3+ HBSS (500 mL HBSS (21-021-CM, Gibco) supplemented with 1% 1 M HEPES (15630080, Gibco), 1% 100 mM sodium pyruvate (25000CI, Corning), 0.5% penicillin-streptomycin, and 100 mL ddH₂O) and stored in the same ice-cold solution. Hippocampus tissues were dissected following euthanasia as previously described^33^, and stored in ice-cold Hibernate E solution (NC0285514, Fisher Scientific) supplemented with 1% B-27 Plus (A3582801, Gibco) and 1% GlutaMAX.

The hippocampus tissues were transferred to a tissue culture hood, washed repeatedly with sterile 3+ HBSS in a 15 mL conical tube, and digested with papain-containing HBSS solution (20 U/mL papain (LS003126, Worthington), 5 mM L-cysteine, 1.1 mM EDTA (pH 8.5)) at 37°C for 7– 10 minutes. DNase I (LS006355, Worthington) was added halfway through digestion. The reaction was quenched with fetal bovine serum (FBS), followed by sequential washes with 3+ HBSS, Neuron Basal Medium (10888022, Gibco) and plating medium to remove residual enzymes.

Tissues were gently dissociated into a single-cell suspension by pipetting in 1 mL plating medium (complete neuronal medium (Neuron Basal Medium supplemented with 2% B-27 Plus, 1% GlutaMAX, and 1% penicillin-streptomycin) with an additional 5% FBS). The suspension was passed through a cell strainer to remove undissociated cells. Cells were counted, diluted in plating medium, and seeded at 7-9 × 10³ cells/well. After approximately 3 hours, once cells had adhered to the well bottom, the plating medium was replaced with complete neuronal medium (without FBS) to prevent the overgrowth of non-neuronal cell types. Neurons were maintained in a humidified incubator at 37°C with 5% CO_2_.

### 2.6 α-syn PFF transmission assay

Cultured mouse primary hippocampus neurons were co-incubated with 10 µM pesticide and α- syn PFFs at 7 days *in vitro* (DIV). The α-syn PFF transmission assay was conducted as previously described^31^. Briefly, α-syn PFFs were diluted in sterile PBS (Corning), sonicated for 10 cycles, then diluted in neuronal media containing 10 µM pesticide with a final DMSO concentration of 0.1%. At seven days post treatment, half of the neuronal medium was replaced with fresh media. Primary neurons were treated for 14 days, fixed with 4% and evaluated for both toxicity and the development of α-syn pathology as previously reported^34^. Briefly, neurons were fixed with 4% paraformaldehyde/4% sucrose in PBS followed by permeabilization with 1% Triton X-100. To stain insoluble pathological synuclein, cells were incubated overnight at 4°C with phospho-α-syn (Ser129) Antibody 81A diluted 1:4000 in blocking buffer. Neuronal viability was assessed simultaneously by staining with neurofilament light chain (NFL) antibody diluted 1:4000 in blocking buffer. Following primary antibody incubation, cells were washed with PBS and incubated for 1 hour at room temperature with secondary antibodies: goat anti-mouse IgG2a 568 and goat anti-rabbit 488, both diluted 1:2000 in blocking buffer, along with 1 µg/mL DAPI for nuclear staining. Cells were washed again with PBS to remove unbound secondary antibodies and DAPI before imaging. Signal intensity and cell count data were collected from the images. Finally, 81A/NFL levels for each condition were normalized to the α-syn PFF condition, and a hit was defined as having an 81A/NFL normalized to α-syn PFF value > 1 in 2 of 3 trials.

### 2.7 LIVE/DEAD assay in SK-N-MC Cells

Human neuroblastoma SK-N-MC cells were grown and maintained at 37°C and 5% CO_2_ in DMEM with 10% FBS, 5% penicillin/streptomycin. To determine the highest nonlethal concentration (>25% increased cell death relative to vehicle), SK-N-MC human neuroblastoma cells were treated with 0.1 µM -30 µM concentrations of each pesticide for 24 hours. Cell viability was determined as a ratio of ethidium homodimer-1 to calcein-AM fluorescent signal as read by microplate reader and normalized to vehicle-treated cells per manufacturer’s instructions for the Live and Dead Cell Assay (ab115347, Abcam, Waltham, MA, USA). The highest nonlethal concentration for most pesticides in the screen was 10 µM - 30 µM, with few exceptions. (Supplemental Figure S1). To standardize the screen, 10 µM was the concentration used unless otherwise noted.

### 2.8 Autophagy assay in live SK-N-MC cells

Human neuroblastoma SK-N-MC cells were grown at 37°C and 5% CO2 in complete DMEM ( 10% FBS, 5% penicillin/streptomycin). SK-N-MC cells were treated with either 0.1% dimethyl sulfoxide (DMSO) or the highest nonlethal concentration of pesticide in 0.1% DMSO for 24 hours at 37°C. Labelling of autophagosomes by Autophagy Detection Kit green detection reagent (ADK; ab129484, Abcam, Cambridge, UK) and lysosomes by Lysotracker Red DND-99 (LTR; L7528, Invitrogen). Briefly, after cells were treated with vehicle or pesticide in complete DMEM for 24 hours, cells were washed with 1X Assay buffer supplemented with 5% FBS. Cells were then incubated with 100 µL of Microscopy Detection Reagent (2µL Green Detection Reagent/ 1 mL 1X Assay buffer, 5% FBS) with 1 µL Lysotracker for 30 minutes at 37°C protected from light. Cells were washed twice with 200 µL 1X Assay buffer, 5% FBS before 100 µL of 1X Assay buffer, 5% FBS was added to each well. SK-N-MC cells were immediately live-imaged using laser scanning confocal microscopy (LSCM, Leica Microsystems Inc, Buffalo Grove, IL, USA); 63x oil-immersion objective with 2x zoom was used to capture images for analysis. Paired controls were included for each experimental well to account for changes in autophagy over the acquisition period.

### 2.9 Autophagosome and lysosome foci image analysis

Dye-labelled autophagosome and lysosome foci and cell counts were measured using ImageJ (National Institutes of Health, Bethesda, MD, USA) image processing software with “FociPicker3D” package^35^. Briefly, vehicle- and positive control-treated samples were used to manually count foci to establish the threshold to accurately identify foci with FociPicker3D. The threshold was set uniformly across all images. Foci counts were normalized to the number of cell outlines counted manually in collapsed z-stack images. Autophagosome and lysosome foci/cell counts were further normalized to the paired vehicle-treated condition and statistical analyses were conducted on the dataset. Pesticides were considered a hit if the pesticide significantly increased or decreased foci/cell counts (p≤0.05) as determined by a two-tailed Student’s T-Test.

### 2.10 Pesticide-wide Association Study (PWAS)

We have previously performed a PWAS epidemiologic screen of 288 pesticides, linking 53 to PD at a false discovery rate (FDR)<0.05 and 68 at FDR<0.10 ^30^. Briefly, we used 1653 participants of the PEG study (n=829 PD patients and n=824 controls) to test each pesticide for association with PD in an untargeted, agnostic manner. PEG is a population-based case-control study set in three agricultural counties in Central California (Kern, Fresno, and Tulare)^36^. For each study participant, we link lifetime geocoded residential and workplace address histories to the California’s pesticide use report (PUR) database, which documents 50 years of agricultural application of hundreds of pesticides. We then estimate ambient exposure to each pesticide based on residential and workplace proximity to commercial pesticide applications, for example due to living near farms applying pesticides. We use a geospatial algorithm which combines the PUR database with maps of land-use and crop cover to determine for each individual pesticide active ingredient in the PUR, the reported pounds of pesticide applied per acre within a 500m buffer around specific locations, such as addresses, yearly since 1974. For each pesticide, in the PUR and each PEG participant, we determined the average pounds of pesticide applied per acre per year within a 500m buffer of each residential and workplace address over the study window (1974 to 10 years prior to index date, which was PD diagnosis for patients or interview date for controls). We then assessed each pesticide individually for PD risk in a pesticide-wide association study (PWAS). More detail has been published ^30^. Here, we assessed each of the 62 pesticides included in this screen individually for PD risk in a PWAS. The results of this PWAS were integrated with output from the two cell-based screens to highlight pesticides linked to PD through multiple levels and select pesticides for the *in vivo* analysis.

### 2.11 Pesticide treatment in ZF

Manually dechorionated 1 dpf *vmat2:*eGFP ZF embryos were incubated with vehicle or pesticide (0.1 µM -10 µM) in E3 buffer [5 mM NaCl, 0.17 mM KCl, 0.33 mM MgSO_4_, 0.33 mM CaCl_2_,]) with a final concentration of 0.1% dimethyl sulfoxide (DMSO)], at 28.5°C for 6 days. ZF were treated at a density of 2 larvae/mL, and typically no more than 15 larvae were treated per well. ZF were regularly monitored during the treatment period for significant morphological abnormalities and stress, and attrition due to treatment.

### 2.12 ZF brain image acquisition and neuron counts

ZF larvae were anesthetized with <0.01% Syncaine (MS-222) (Syndel, Ferndale, WA, USA) and fixed in 4% PFA. Larvae were washed in PBS, antibody labeled, and cleared in 100% glycerol before ZF whole brain tissues were dissected and mounted onto coverslips for confocal imaging at 40x magnification on a Leica SPE. Briefly, larvae were incubated with peroxide (3% H2O2, 0.8% KOH in PBS) for 8 minutes to clear pigment, permeabilized with Proteinase K at 10 µg/mL for 12 minutes and blocked in 5% lamb serum/5% donkey serum in 0.1% Triton X-100 in PBS. Neurons were immunolabeled with primary monoclonal chicken anti-GFP antibodies (1:1000, A10262, Thermo Fisher Scientific), detected with goat anti-chicken Alexa Fluor 488 (1:1000, A11039, Thermo Fisher Scientific). Larvae were then washed and cleared in 100% glycerol. LSCM was used to image ZF larvae (7 dpf); 40x-oil immersion objective with 1x zoom was used to capture the telencephalic and diencephalic brain regions for analysis. Alexa Fluor 488+ neurons in *vmat2*:eGFP ZF were counted using ImageJ (NIH). Images were randomized and Alexa Fluor 488+ positive neurons were counted in the telencephalic and diencephalic regions. Once the data set was counted, the images were unblinded and the data were collated and analyzed.

### 2.13 Statistical analysis

Primary neuron and ZF aminergic neuron data are represented as mean ± SEM. Foci counts are represented as log_2_ fold-change from vehicle-treated cells. Sample size (N) are provided per condition in each figure legend. Pair-wise comparisons for foci counts in SK-N-MCs were analyzed using Student’s two-tailed t-test. Dose response to pesticide in ZF was analyzed by one-way ANOVA with Tukey’s post-hoc comparisons test. Statistical significance was considered at P < 0.05; groups being compared are identified in the figure legend. Statistical analyses were conducted using GraphPad Prism v. 9.5.1 software (GraphPad, Boston, MA, USA).

## 3. Results

### 3.1 Identification of pesticides that promote α-syn PFF transmission

The 62 pesticides were evaluated for their ability to promote pathogenic α-syn transmission in primary mouse neurons. Primary cultures were treated for 14 days with α-syn PFFs or pesticide and α-syn PFFs, fixed, and the amount of α-syn pathology was measured by indirect immunofluorescence with antibody against phosphorylated α-syn (p-α-syn), NFL, and NeuN (Figure 2 A-B). Most of the pesticides did not exhibit increased toxicity at 10 µM compared to the PFF-only condition as measured by NFL labelling (Figure 2C, Supplement Figure S2A). Twenty-nine pesticides showed an increased amount of p-α-syn/NFL levels compared to the PFF-only condition (Figure 2D, Supplement Figure S2B), suggesting that these pesticides promote pathological α-syn transmission. The largest increases in pathological α-syn were found after exposure to benomyl, captan, acephate, abamectin, 2,4-D and bromacil in the presence of α-syn PFFs. Results for NFL and p-α-syn/NFL levels for the full panel of pesticides in the screen are outlined in Supplemental Figure S2.

**Figure 2.**
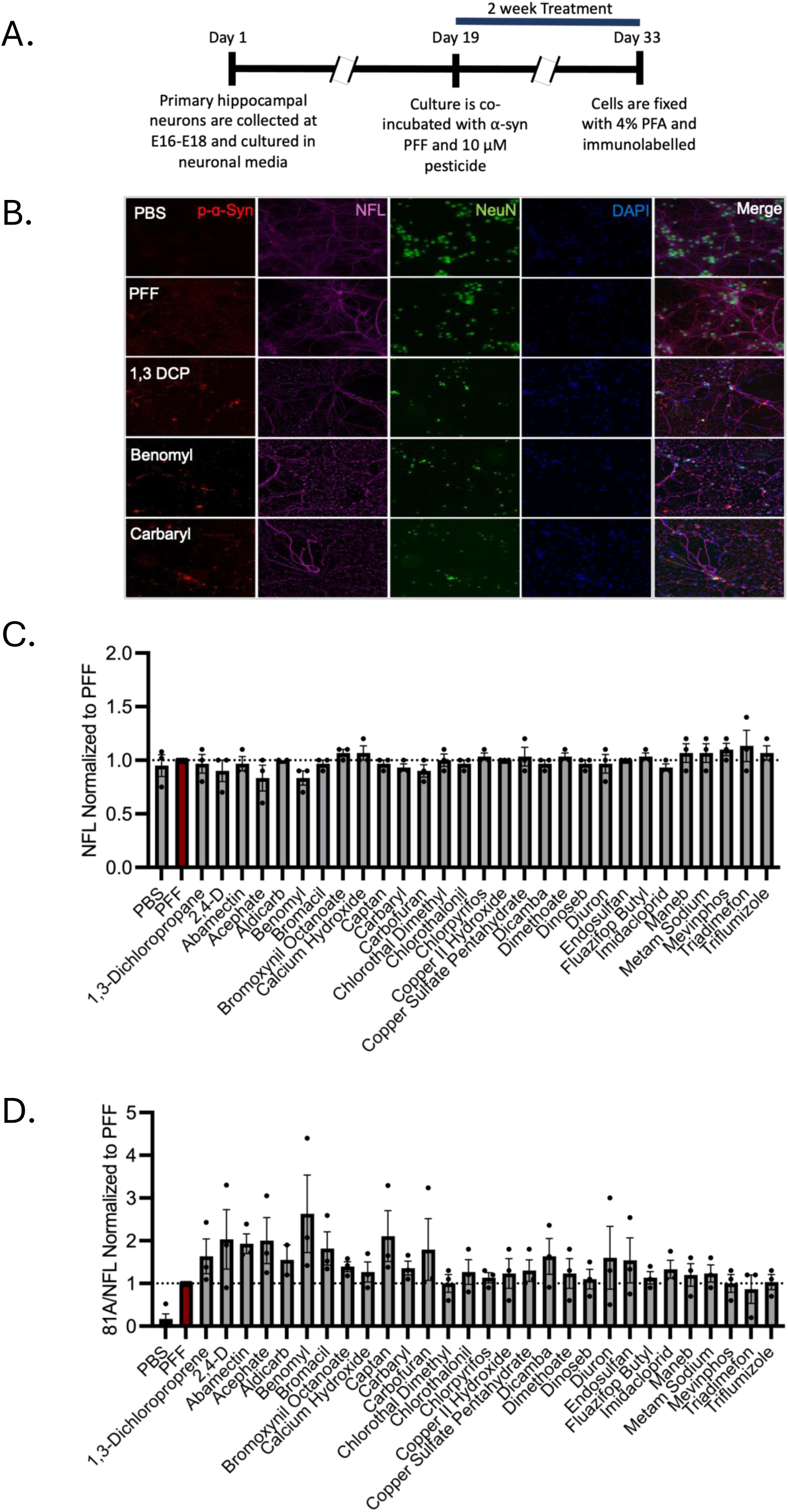
Pesticide-induced transmission of pathogenic alpha-synuclein pre-formed fibrils (ɑ-Syn PFFs) in mouse primary hippocampal neurons. **A)** Treatment schema outlining the treatment timeline from collection to co-incubation to fixation and immunolabelling of primary cultures. **B)** Primary cultures were treated for 14 days with the vehicle or pesticide and co- incubated with ɑ-syn PFFs, fixed, and immunolabelled with phosphorylated ɑ-syn (red, anti-p-ɑ- syn), neurofilament light (magenta, anti-NFL), NeuN (green, anti-NeuN), and DAPI (blue, nuclear stain). **C)** Pesticides were found to not have exhibit increased neuron loss relative to the PFF only condition as shown by NFL levels. **D)** Twenty-eight pesticides show increased levels of p-ɑ- syn/NFL levels compared to PFF only condition. Images shown are representative images for each condition. Data shown as mean ± SEM and is normalized to PFF only condition; N=3 experimental replicates. The pesticide is considered a hit when p-ɑ-syn/NFL > 1 in 2 of 3 independent trials. Abbreviations: NFL, neurofilament light chain; ɑ-syn, alpha-synuclein; PFF, pre-formed fibrils; 1,3-DCP, 1,3-dichloroproprene.

### 3.2 Identification of pesticides that alter autophagy in SK-N-MC cells

All 62 pesticides were evaluated for their effects on autophagy after 24-hour incubation in SK-N-MC cells (Figure 3A). For this screen, we used the highest nonlethal concentration for each chemical. The cytotoxicity of the panel of pesticides in SK-N-MC cells was assessed using a LIVE/DEAD dye assay with concentrations ranging from 0.1 µM – 30 µM. The treatment was well- tolerated up to 30 µM for most of the pesticides, evidenced by less than 25% cell death relative to control (Supplemental Figure S1). The 10 µM concentration was used for the autophagy assay in live SK-N-MC cells, except 1 µM was used for rotenone and chlorothalonil due their toxicity at higher concentrations. Treatment with the autophagy inhibitor, chloroquine, was used as a positive control to show inhibition of autophagy demonstrated by the increase in autophagosome and lysosome puncta assessed by confocal microscopy (Figure 3B). The screen revealed 16 pesticides that significantly altered the number of autophagosomes and 14 pesticides that significantly altered the number of lysosomes in SK-N-MC cells relative to vehicle-treated cells for a total of 22 pesticide hits (Figure 3C). While none of the pesticides achieved the high levels of both autophagosomes and lysosomes present with the autophagy inhibitor, ziram, chlorothal dimethyl, napropamide, and prometryn all exhibited the largest increases in autophagosome and lysosome foci relative to vehicle (Figure 3C). There was a subset of chemicals that behaved in a similar pattern of moderate increases in both autophagosome and lysosome labeling (copper hydroxide, diuron, fenarimol, mancozeb, and triflumizole) (Figure 3C). There was also a subset of pesticides that exhibited a reduction in autophagosome and lysosome labeling (benomyl, carbofuran, rotenone, sulfur, and zineb) (Figure 3C). Results for the full panel of screened pesticides through the autophagy assay can be found in Supplemental Figure S3 and Supplement Tables S2 and S3.

**Figure 3.**
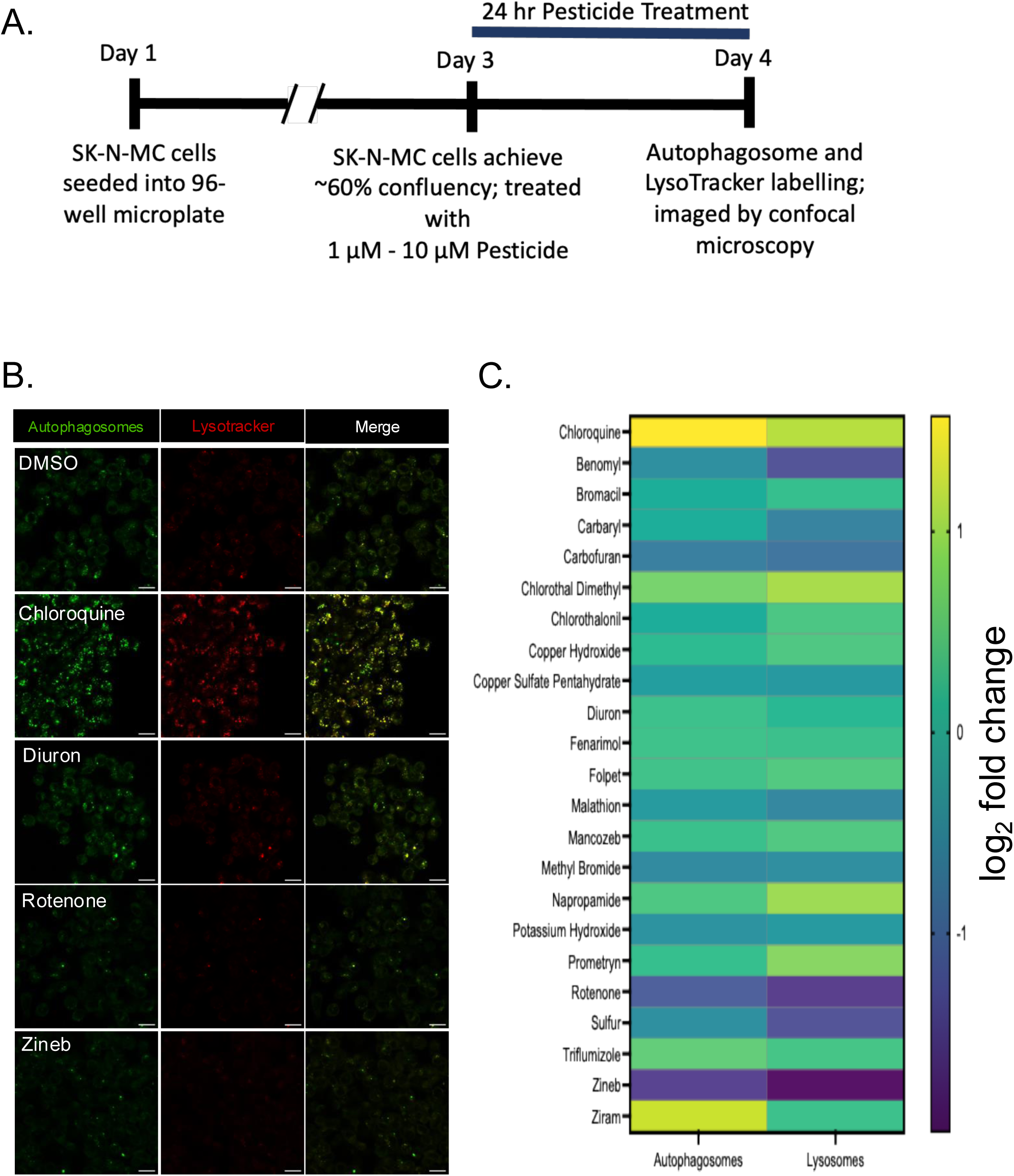
Screen for changes in autophagy components, autophagosomes and lysosomes, after pesticide exposure in SK-N-MC cells. **A)** Treatment schema outlining the treatment timeline from plate seeding to treatment to fixation and dye-labelling. **B)** SK-N-MCs were treated pesticide for 24hr with the vehicle or pesticide, fixed, and rapidly labelled with Autophagy Dye (green, Autophagy Detection Reagent) and Lysotracker (red, Lysotracker Red DND-99) for immediate live-cell imaging. Images shown are representative images for each condition. **C)** Twenty-two pesticides show significant perturbations (increased or decreased foci counts) to autophagosome and lysosome pools relative to vehicle and are considered hits. Data was analyzed as mean ± SEM and represented as log2 fold change from vehicle (0.1% DMSO); N= 3 images/well, 2 biological replicates with paired control wells. Pair-wise comparisons were conducted by Student’s T-test; raw foci counts for each pesticide was compared to a paired- vehicle condition. The pesticide is considered a hit at P<0.05 for either the autophagosome or lysosome category.

### 3.3 Pesticides with an increased odds ratio (OR) for PD in a case-control study of PD

The third component of the screen involved utilizing the results from the untargeted screening of individual pesticides for association with PD (PWAS). The analysis tested 288 pesticides individually for association with PD in 1653 study participants, controlling for age, sex, race/ethnicity, education level, and index year (of diagnosis or interview) to account for temporal trends in pesticide use. Exposure was estimated at both the participant’s residence and workplace and associations were independently evaluated for each exposure location. Overall, 68 pesticides were linked to PD (25 at FDR<0.01, 28 at 0.01≤FDR≤0.05, and 15 at 0.05<FDR<0.10)^30^. Of the 62 pesticides screened in the cell assays, 34 had an association with PD, with sodium chlorate, kelthane (dicofol), and prometryn showing the largest OR for PD.

### 3.4 Triple-hits assessed for inducing aminergic neuron loss in ZF

We assumed that the chemicals that were positive in all three screens would be the most likely to be causatively associated with increased PD risk. Thus, pesticides that 1) promoted α- syn PFF transmission, 2) significantly altered either autophagosome or lysosome foci counts, and 3) were found to have an association with PD in our PWAS were defined as a positive hit. While each of the screens resulted in several hits, only 6 were positive in all three screens (i.e., triple hits) (Figure 4, Table 1). To further test the triple hits for their ability to contribute to some of the pathology seen in patients, we utilized ZF as a model organism. ZF has emerged as a useful animal model to study neurodegenerative processes due to their optical clarity, rapid development, and conserved brain structures with mammals^37^. The 6 pesticides that met the triple-hit criteria (carbaryl, carbofuran, copper sulfate, chlorthal dimethyl, copper hydroxide, and diuron) were evaluated for aminergic neuron (*VMAT2*+) toxicity. Significant *VMAT2*+ neuron loss (P<0.05) was observed with carbofuran (at 10 µM) and copper hydroxide (at 10 µM), carbaryl (at 1 µM), and copper sulfate (at 0.1 µM) in the telencephalon of 7 dpf ZF (Figure 5A-G). Further, a significant loss of *VMAT2*+ neurons (P<0.05) was also observed in the diencephalon region with carbofuran (at 1 µM), chlorthal dimethyl (at only 1 µM), and copper sulfate (at 0.1 µM) in 7 dpf ZF (Figure 5A-G).

**Figure 4.**
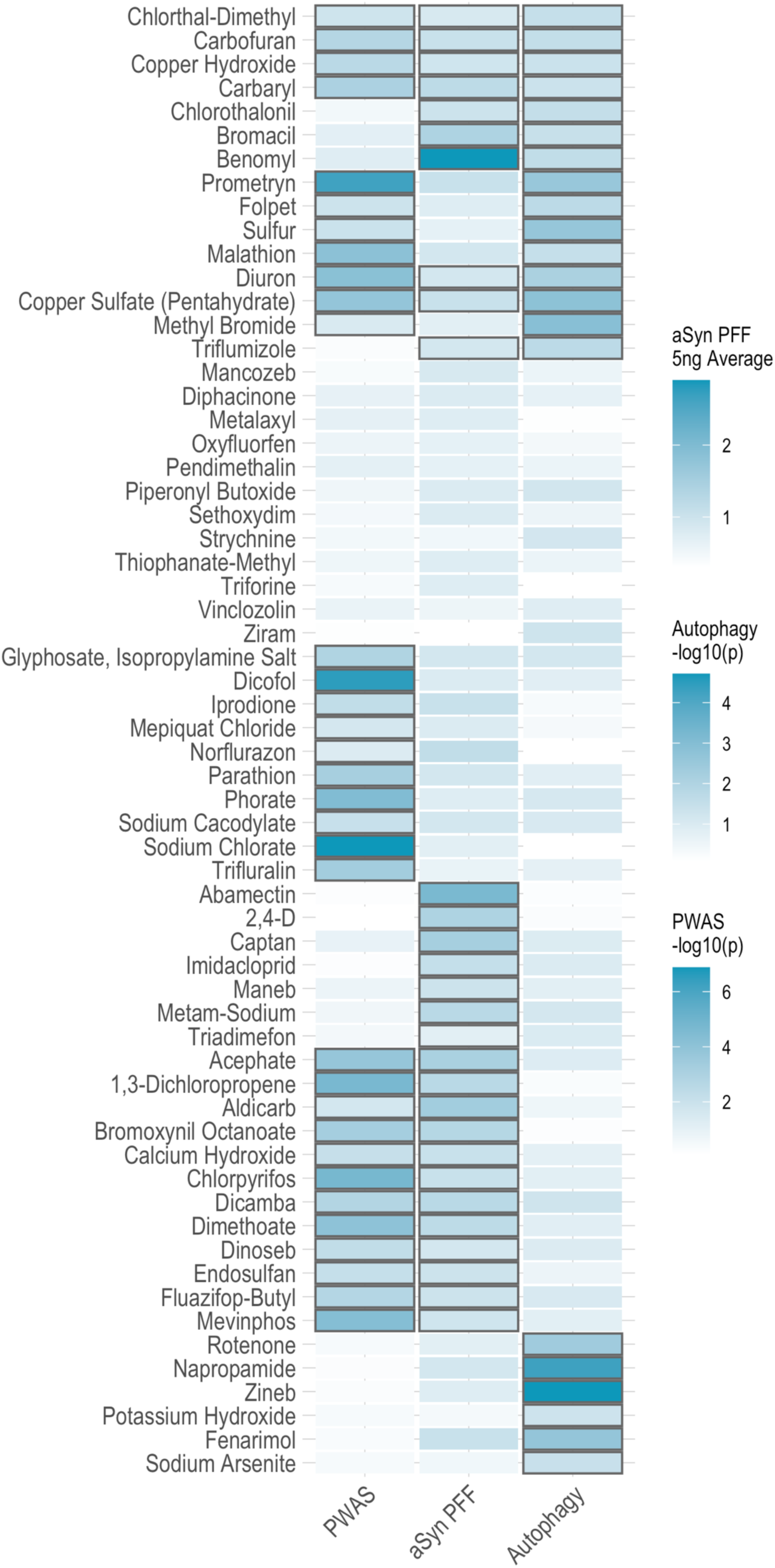
Six pesticides are identified as hits across all three screens. Carbaryl, carbofuran, copper sulfate, chlorthal dimethyl, copper hydroxide and diuron are identified as triple hits in the independent screens.

**Table 1.** Multiscreen hits show alterations to autophagosome and lysosome foci counts relative to vehicle in SK-N-MC cells.

|  |  | Vehicle |  |  |  | Autophagosomes |  |  |  |  |
| --- | --- | --- | --- | --- | --- | --- | --- | --- | --- | --- |
| Concentration | Pesticide | Mean | ± | SEM | N | Mean | ± | SEM | P-Value | N |
| 10 µM | Chloroquine | 1 | ± | 0.171438017 | 5 | 2.975271815 | ± | 0.258205537 | 0.0003 | 4 |
| 10 µM | Carbaryl | 1 | ± | 0.140412554 | 3 | 1.016411881 | ± | 0.076909711 | 0.91635 | 4 |
| 10 µM | Carbofuran | 1 | ± | 0.077149242 | 3 | 0.59852322 | ± | 0.097426095 | 0.02879 | 4 |
| 10 µM | Chlorothal Dimethyl | 1 | ± | 0.270604265 | 3 | 1.687698181 | ± | 0.080940347 | 0.03811 | 4 |
| 10 µM | Copper Hydroxide | 1 | ± | 0.061945839 | 3 | 1.203503173 | ± | 0.102975449 | 0.23358 | 6 |
| 10 µM | Copper Sulfate | 1 | ± | 0.026531948 | 3 | 0.838436913 | ± | 0.01149274 | 0.00159 | 4 |
| 10 µM | Diuron | 1 | ± | 0.048364665 | 3 | 1.303864584 | ± | 0.048943999 | 0.00771 | 4 |
|  |  | Vehicle |  |  |  | Lysosomes |  |  |  |  |
| Concentration | Pesticide | Mean | ± | SEM | N | Mean | ± | SEM | P-Value | N |
| 10 µM | Chloroquine | 1 | ± | 0.233694148 | 5 | 2.178084956 | ± | 0.322723174 | 0.01895 | 4 |
| 10 µM | Carbaryl | 1 | ± | 0.084540268 | 3 | 0.621155856 | ± | 0.104054687 | 0.04445 | 4 |
| 10 µM | Carbofuran | 1 | ± | 0.152068902 | 3 | 0.540764712 | ± | 0.104148527 | 0.04868 | 4 |
| 10 µM | Chlorothal Dimethyl | 1 | ± | 0.263581856 | 3 | 2.058851747 | ± | 0.241676435 | 0.03255 | 4 |
| 10 µM | Copper Hydroxide | 1 | ± | 0.198055306 | 3 | 1.423333514 | ± | 0.075852827 | 0.04222 | 6 |
| 10 µM | Copper Sulfate | 1 | ± | 0.08323006 | 3 | 0.795621614 | ± | 0.089988183 | 0.16909 | 4 |
| 10 µM | Diuron | 1 | ± | 0.073485843 | 3 | 1.174974755 | ± | 0.107197543 | 0.26953 | 4 |

**Figure 5.**
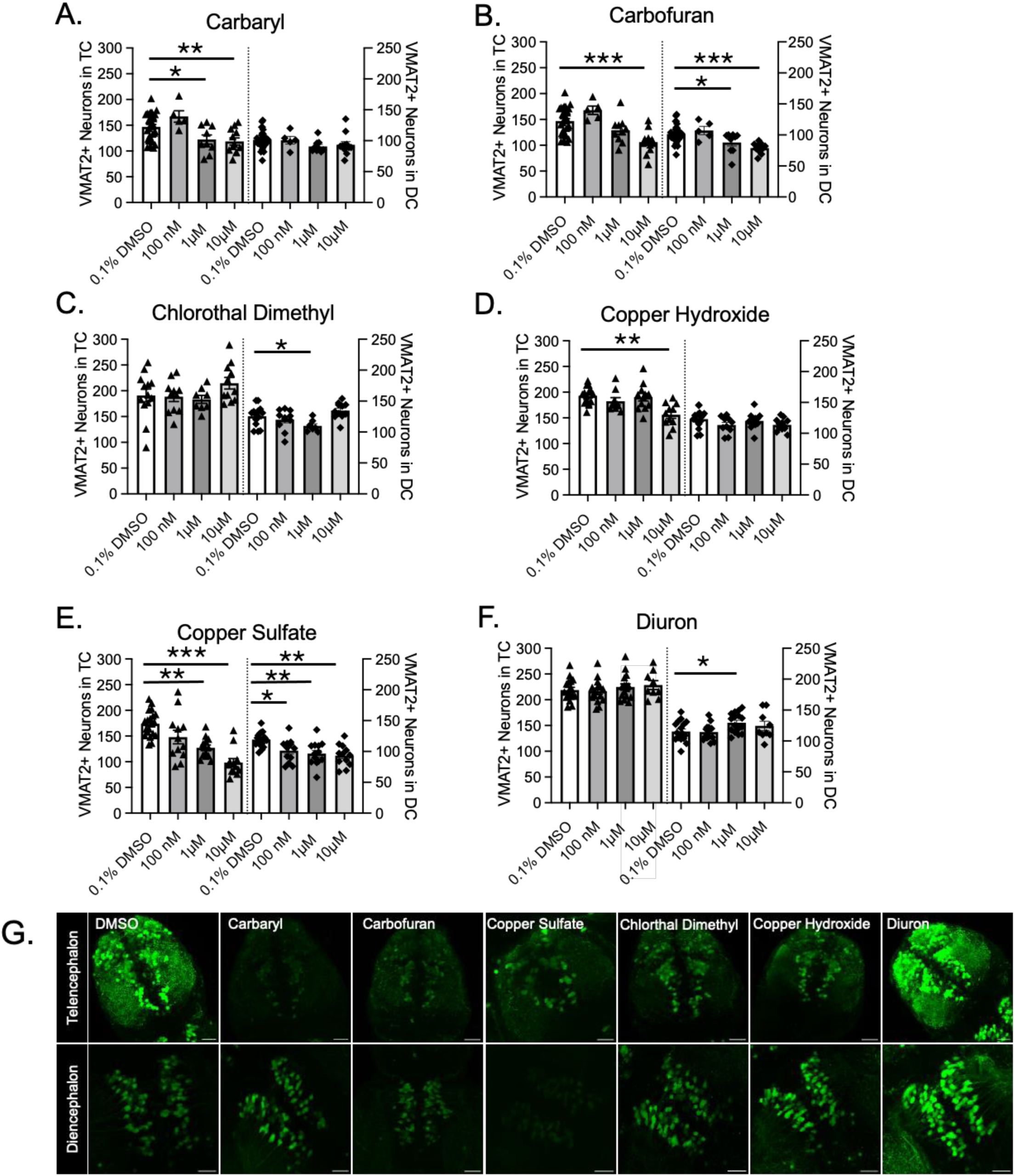
Carbaryl, carbofuran, copper hydroxide, and CSP induce aminergic neuron loss in 7dpf zebrafish brain. The number of *VMAT2+* neurons in the ZF telencephalon and diencephalon was evaluated after pesticide exposure at multiple doses after a 6d exposure period. The lowest dose at which we observed a significant reduction in the number of neurons is identified in each panel. **A)**. Carbaryl reduced the number of *VMAT2+* neurons in the telencephalon (at 1 µM), and showed no effect in the diencephalon. **B)** Carbofuran reduced the number of *VMAT2+* neurons in both the telencephalon and diencephalon (at 10 µM and 1 µM, respectively). **C)** Chlorthal dimethyl reduced the number of *VMAT2+* neurons in the diencephalon (at 1 µM) telencephalon but showed no effect in the telencephalon. **D)** Copper hydroxide reduced the number of *VMAT2+* neurons in the telencephalon (at 10 µM), and showed no effect in the diencephalon. **E)** Copper sulfate reduced the number of *VMAT2+* neurons in both the telencephalon and diencephalon (at 0.1 µM). **F)** Diuron did not alter the number of *VMAT2+* neurons in the telencephalon, but showed a modest increase at only 1 µM in the telencephalon. **G)** Representative images of ZF telencephalon and diencephalon brain regions for each condition at the highest concentration of pesticide evaluated (10 µM). Data shown as mean ± SEM; N= 5-14/condition. Dose response comparisons were conducted by one-way ANOVA with Dunnett’s post-hoc multiple comparisons test; all doses of pesticide were compared to vehicle (0.1% DMSO) alone. *p≤0.05, **p≤0.01, ***p≤0.001

## 4. Discussion

The association of pesticides with incident PD has been well established by many studies, but very few individual chemicals that confer this risk have been described. Identification of individual toxins is essential to determining whether the association is causative. There is no single accepted method to determine causality of associations, but the combination of epidemiological findings with cellular and animal studies has been employed to establish not only biological plausibility but also allow for causal inference. Here, we identified 4 pesticides (carbaryl, carbofuran, copper hydroxide, copper sulfate) that altered 2 adverse outcome pathways (AOPs) in PD (i.e., promote a-syn transmission and alter autophagy) and also reduced aminergic neuronal populations in an *in vivo* model. Using a mechanistic approach, we have identified 4 pesticides that have not been previously implicated in PD risk beyond our epidemiologic studies and induce molecular changes consistent with disease. These chemicals thus were identified not only as environmental risk factors for developing PD but exhibit considerable potential to alter AOPs critical to disease initiation and propagation. This study provides biological plausibility for the association between pesticide exposures and disease risk and also highlights critical pathways that may link environmental exposures to disease processes. The findings of this study offer a strong basis for further investigation in additional mammalian models and epidemiological studies of PD.

As PD pathophysiology typically develops over decades in humans, an AOP approach is appropriate for identifying causative environmental toxins. Since disruption of autophagy and α- syn transmission can lead to the development of PD over several years, we believe using these 2 AOPs as screening tools is a strength and adds strong biological plausibility for these 4 pesticides and support causal inference. The use of the PEG cohort and the PWAS approach for identifying PD-associated pesticides is also a strength. Few cohorts have the ability to estimate subjects’ exposures to individual chemicals for over 45 years and not rely on subject recall for exposure assessment. Furthermore, the diagnosis of PD was made by a movement disorders specialist who followed them over time to further confirm an accurate diagnosis.

Despite all of these strengths, we acknowledge there are limitations to this approach. It is likely that not all patients get PD due to altered autophagy and the a-syn spread theory of progression is not universally accepted. Although we likely have missed some pesticides that contribute to risk since there are other relevant AOPs in PD (e.g., mitochondrial dysfunction) that we did not test for and may be able to alter risk. With that said, we chose to improve our chances of identifying some causative agents by using stringent criteria in our studies (i.e., positive in all 3 screens). Another limitation of our study is the uncertainty of whether the selected concentrations are relevant to human exposures. This is extremely difficult to determine but we carefully considered actual use concentrations for these chemicals to maintain relevant human exposure levels. For example, based on the manufacturer’s instructions, 2 grams of the copper fungicide should be applied to 1 square meter which would be estimated to be in the low millimolar range on the surface (Southern Ag - Liquid Copper Fungicide). In the case of copper sulfate, use is controlled for various purposes and can range from 0.25 ppm to 10 ppm on crops and in water which corresponds to a low micromolar range (approximately 1 - 62 µM) (US EPA Reg. No. 88633-3). Thus, we focused our evaluation on low micromolar concentrations in our screens and observed reductions in *VMAT2+* neurons as low as 0.1 µM (copper sulfate) in our ZF model which are likely environmentally relevant concentrations.

ZF have several advantages given that they develop rapidly with well-formed aminergic neurons, including dopaminergic neurons, in 3 days. They are readily genetically modified to express fluorescent reporters, are transparent allowing for easy imaging of intact larvae, and ZF share a high homology with most mammalian genes. The means of ZF exposures is a potential limitation as the ZF were exposed whole-body in a water medium with pesticide. However, we suggest that the whole-body exposure accounts for the three primary routes of exposure (inhalation, ingestion, and dermal) and may provide useful insights for pesticide prioritization ahead of exposure route-specific studies. Another potential limitation is the use of developing ZF in a disease of the aged. The small size of developing ZF allows us to treat in a multi-well format and leverage the model to quickly and effectively screen for pesticide-induced neuronal toxicity^38^. With these findings, we offer 4 prioritized triple-hit pesticides for in-depth mechanistic investigation, in addition to the hits for each individual *in vitro* screen.

In summary, we developed a multiscreen mechanistic approach to identify pesticides that increase the risk of developing PD and alter pathways critical to disease pathogenesis. We identified 4 pesticides that act to alter autophagy and α-syn transmission, are associated with PD in our epidemiologic analysis, and induce aminergic neuron loss in a ZF model. The results of this study offer a strong basis for further mechanistic studies of these pesticides in other PD experimental models to establish causality and develop potential treatments.

## Acknowledgements

This study was supported by the National Institutes of Health Environmental Health Sciences (NIEHS) T32ES015457 (MA), R01ES031106 (BR), and The Levine Foundation (JMB).

## Author Contributions

Conceptualization: MA, LMB, JMB

Methodology: MA, CP, KCP

Investigation: MA, LMB, AMK, KMMH, SL

Visualization: MA, KCP

Supervision: JMB, KCP, CP, BR

Funding acquisition: BR, JMB

Project administration: JMB

Writing – original draft: MA, JMB

Writing – review and editing: MA, LMB, AMK, KMMH, SL, CP, KCP, BR, JMB

## Data Availability Statement

All data are included in the manuscript and may be made available upon request.

---

**Supplement Table S1.** Pesticide information

| Supplement Table S1. Pesticide information |  |  |
| --- | --- | --- |
| Pesticide | Vendor | Catalog Number |
| 1,3-DICHLOROPROPENE | Chem Service Inc | N-10193-500MG |
| 2-4-D | Sigma-Aldrich | 31518-250MG |
| ABAMECTIN | Chem Service Inc | N-10995-100MG |
| ACEPHATE | Sigma-Aldrich | 11-101-9093 |
| ALDICARB | Sigma-Aldrich | 33386-100mg |
| BENOMYL | Chem Service Inc | N-11138-100MG |
| BROMACIL | Chem Service Inc | N-11330-250MG |
| BROMOXYNIL OCTANOATE | Sigma-Aldrich | 11-101-9958 |
| CALCIUM HYDROXIDE | Sigma-Aldrich | 239232 |
| CAPTAN | Chem Service Inc | N-11400-250MG |
| CARBARYL | Sigma-Aldrich | 11-102-0173 |
| CARBOFURAN | Sigma-Aldrich | 11-102-0498 |
| CHLOROTHALONIL | Chem Service Inc | N-11454-250MG |
| CHLORPYRIFOS | Chem Service Inc | N-11459-100MG |
| CHLORTHAL-DIMETHYL | Chem Service Inc | N-11462-250MG |
| COPPER HYDROXIDE | Chem Service Inc | NG-I7032-1G |
| CUPRIC SULFATE PENTAHYDRATE | Sigma-Aldrich | C-7631 |
| DICAMBA, DIMETHYLAMINE SALT | Chem Service Inc | N-11656-250MG |
| DIMETHOATE | Sigma-Aldrich | 45449-100MG |
| DINOSEB | Chem Service Inc | N-11786-100MG |
| DIPHACINONE | Chem Service Inc | N-11793-100MG |
| DIURON | Sigma-Aldrich | 11-102-0532 |
| ENDOSULFAN | Chem Service Inc | N-10992-250MG |
| FENARIMOL | Chem Service Inc | N-11948-100MG |
| FLUAZIFOP-BUTYL | Chem Service Inc | N-11978-250MG |
| FOLPET | Sigma-Aldrich | 32057-250MG |
| GLYPHOSATE, ISOPROPYLAMINE SALT | Chem Service Inc | N-12134-250MG |
| IMIDACLOPRID | Chem Service Inc | N-12206-100MG |
| IPRODIONE | Chem Service Inc | N-12220-100MG |
| KELTHANE (DICOFOL) | Chem Service Inc | N-12290-100MG |
| MALATHION | Sigma-Aldrich | 11-102-0185 |
| MANCOZEB | Sigma-Aldrich | 45553-250MG |
| MANEB | Chem Service Inc | N-12355-250MG |
| MEPIQUAT CHLORIDE | Chem Service Inc | N-12371-100MG |
| METALAXYL | Chem Service Inc | N-12380-100MG |
| METAM SODIUM | Sigma-Aldrich | 45570-250MG |
| METHYL BROMIDE | Chem Service Inc | S-12417M8-1ML |
| MEVINPHOS | Chem Service Inc | NC1440273 |
| NAPROPAMIDE | Chem Service Inc | N-11585-250MG |
| NORFLURAZON | Chem Service Inc | N-12668-100MG |
| OXYFLUORFEN | Chem Service Inc | N-12742-100MG |
| PARATHION METHYL | Sigma-Aldrich | 11-101-9734 |
| PENDIMETHALIN | Sigma-Aldrich | 36191-100MG |
| PHORATE | Sigma-Aldrich | 11-101-7273 |
| PIPERONYL BUTOXIDE | Chem Service Inc | NC9954965 |
| POTASSIUM HYDROXIDE | Chem Service Inc | NG-I103-1G |
| PROMETRYN | Sigma-Aldrich | 11-101-9387 |
| ROTENONE | Sigma-Aldrich | R8875-1G |
| SETHOXYDIM | Chem Service Inc | N-13210-10MG |
| SODIUM ARSENITE | Chem Service Inc | NG-I121-1G |
| SODIUM CACODYLATE | Thermo Scientific Chemicals | AC214970100 |
| SODIUM CHLORATE | Thermo Scientific Chemicals | AC446411000 |
| STRYCHNINE | Chem Service Inc | N-13231-100MG |
| SULFUR | Sigma-Aldrich | 36576-250MG |
| THIOPHANATE METHYL | Sigma-Aldrich | 45688-250MG |
| TRIADIMEFON | Chem Service Inc | N-13636-500MG |
| TRIFLUMIZOLE | Sigma-Aldrich | 32611-100MG |
| TRIFLURALIN | Sigma-Aldrich | 45700-250MG |
| TRIFORINE | Chem Service Inc | N-13691-250MG |
| VINCLOZOLIN | Sigma-Aldrich | 45705-250MG |
| ZINEB | Sigma-Aldrich | 45707-250MG |
| ZIRAM | Chem Service Inc | N-13761-250MG |

**Supplement Figure 1.**
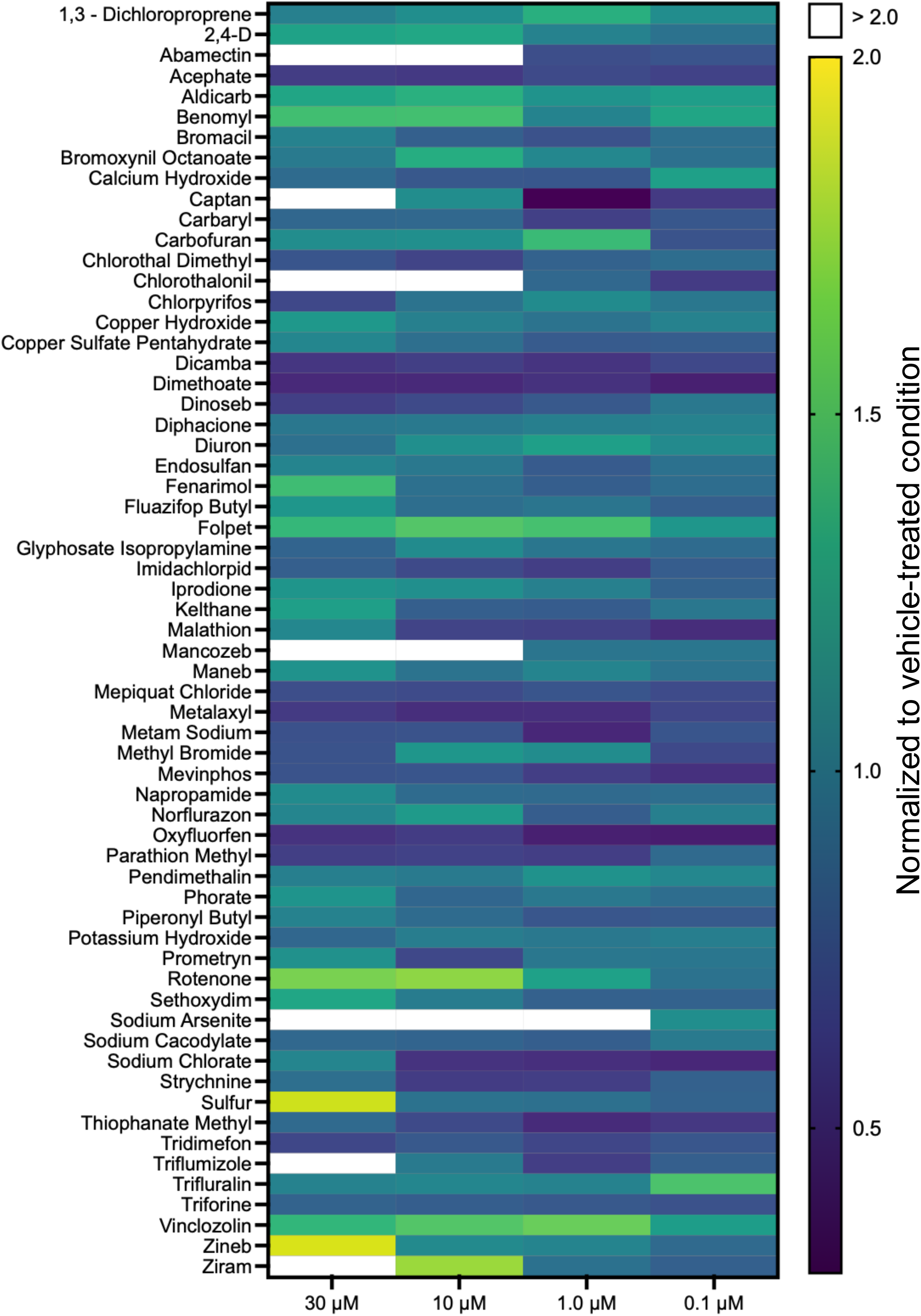
Dose response evaluation of pesticides on cell viability in human neuroblastoma SK-N-MC cells. Dose response studies were conducted to evaluate pesticide- induced lethality. Most pesticides were found to be well-tolerated by SK-N-MC cells at 1µM - 10µM.

**Supplement Figure 2.**
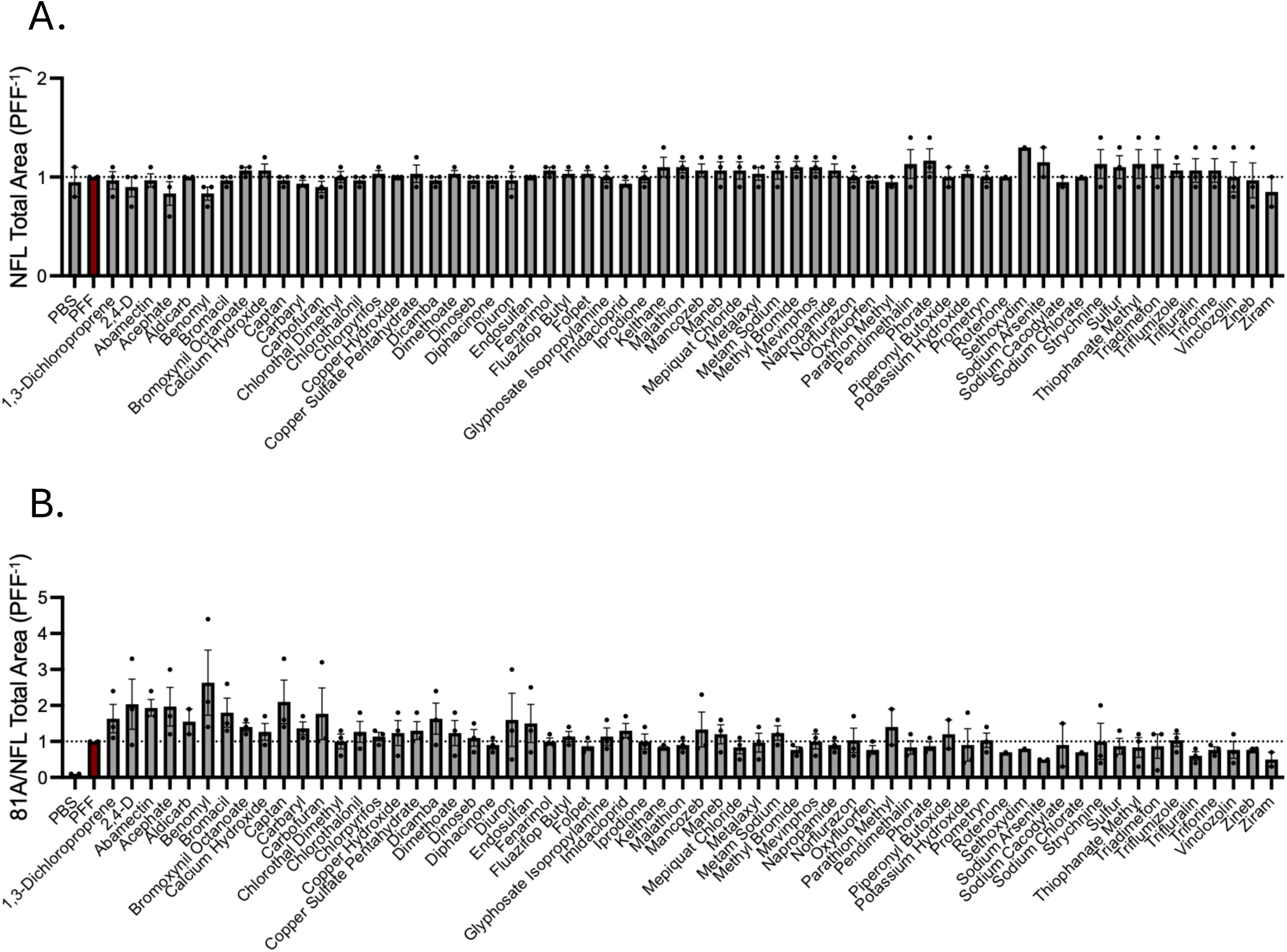
Effects of pesticides on transmission of pathogenic alpha-synuclein pre-formed fibrils in mouse primary hippocampal neurons. **A)** Neuron viability was not affected by most pesticides and PFF treatment evaluated in the screen at 10 µM relative to the α-syn PFF only condition as shown by NFL levels. **B)** Primary cultures treated with pesticides and α-syn PFFs were evaluated for p-ɑ-syn immunofluorescence levels normalized to NFL levels and compared to PFF only condition.

**Supplement Table 2.** Results of autophagosome foci differences in SK-N-MC autophagy assay. Pair-wise comparisons were conducted by Student’s T-test; raw foci counts for each pesticide was compared to a paired-vehicle condition. The pesticide is considered a hit at P<0.05.

| Concentration | Pesticide | Vehicle |  |  | N | Autophagosomes |  |  | P-Value | N |
| --- | --- | --- | --- | --- | --- | --- | --- | --- | --- | --- |
|  |  | Mean | ± | SEM |  | Mean | ± | SEM |  |  |
| 10 µM | 1,3-Dichloropropene | 1 | ± | 0.01705 | 3 | 0.92388 | ± | 0.11298 | 0.59574 | 4 |
| 10 µM | 2,4-D | 1 | ± | 0.11424 | 3 | 1.05754 | ± | 0.07817 | 0.6833 | 4 |
| 10 µM | Abamectin | 1 | ± | 0.11394 | 3 | 0.95371 | ± | 0.04555 | 0.65763 | 6 |
| 10 µM | Acephate | 1 | ± | 0.12641 | 4 | 0.83334 | ± | 0.06815 | 0.24009 | 6 |
| 10 µM | Aldicarb | 1 | ± | 0.18176 | 3 | 1.2354 | ± | 0.12823 | 0.32313 | 4 |
| 10 µM | Atrazine | 1 | ± | 0.15798 | 4 | 1.03474 | ± | 0.18761 | 0.892 | 4 |
| 10 µM | Benomyl | 1 | ± | 0.09751 | 3 | 0.71543 | ± | 0.04932 | 0.03638 | 4 |
| 10 µM | Bromacil | 1 | ± | 0.01509 | 3 | 1.03275 | ± | 0.03335 | 0.50154 | 5 |
| 10 µM | Bromoxynil Octanoate | 1 | ± | 0.10772 | 3 | 1.00834 | ± | 0.03938 | 0.93767 | 4 |
| 10 µM | Calcium Hydroxide | 1 | ± | 0.22647 | 3 | 1.11546 | ± | 0.00797 | 0.56938 | 4 |
| 10 µM | Captan | 1 | ± | 0.10541 | 4 | 1.11368 | ± | 0.08024 | 0.41021 | 5 |
| 10 µM | Carbaryl | 1 | ± | 0.14041 | 3 | 1.01641 | ± | 0.07691 | 0.91635 | 4 |
| 10 µM | Carbofuran | 1 | ± | 0.07715 | 3 | 0.59852 | ± | 0.09743 | 0.02879 | 4 |
| 10 µM | Chloroquine | 1 | ± | 0.17144 | 5 | 2.97527 | ± | 0.25821 | 0.0003 | 4 |
| 10 µM | Chlorothal Dimethyl | 1 | ± | 0.2706 | 3 | 1.6877 | ± | 0.08094 | 0.03811 | 4 |
| 1 µM | Chlorothalonil | 1 | ± | 0.05621 | 3 | 1.00676 | ± | 0.06586 | 0.94668 | 5 |
| 10 µM | Chlorpyrifos | 1 | ± | 0.09953 | 4 | 1.29923 | ± | 0.1607 | 0.16451 | 4 |
| 10 µM | Copper Hydroxide | 1 | ± | 0.06195 | 3 | 1.2035 | ± | 0.10298 | 0.23358 | 6 |
| 10 µM | Copper Sulfate | 1 | ± | 0.02653 | 3 | 0.83844 | ± | 0.01149 | 0.00159 | 4 |
| 10 µM | Dicamba | 1 | ± | 0.17024 | 3 | 1.29636 | ± | 0.0905 | 0.13065 | 6 |
| 10 µM | Dimethoate | 1 | ± | 0.04323 | 3 | 1.1653 | ± | 0.07649 | 0.15063 | 4 |
| 10 µM | Dinoseb | 1 | ± | 0.19952 | 3 | 1.34154 | ± | 0.08595 | 0.115 | 5 |
| 10 µM | Diphacione | 1 | ± | 0.08921 | 3 | 0.76278 | ± | 0.11523 | 0.20499 | 5 |
| 10 µM | Diuron | 1 | ± | 0.04836 | 3 | 1.30386 | ± | 0.04894 | 0.00771 | 4 |
| 10 µM | Endosulfan | 1 | ± | 0.20688 | 4 | 0.75128 | ± | 0.03394 | 0.28031 | 4 |
| 10 µM | Fenarimol | 1 | ± | 0.03994 | 3 | 1.31646 | ± | 0.03741 | 0.00231 | 4 |
| 10 µM | Ferbam | 1 | ± | 0.05835 | 3 | 1.37194 | ± | 0.13454 | 0.0758 | 4 |
| 10 µM | Fluazifop Butyl | 1 | ± | 0.0999 | 3 | 1.10076 | ± | 0.03707 | 0.33461 | 4 |
| 10 µM | Folpet | 1 | ± | 0.06939 | 3 | 1.3237 | ± | 0.06498 | 0.02011 | 4 |
| 10 µM | Glyphosate Isopropylamine | 1 | ± | 0.14767 | 3 | 0.6955 | ± | 0.06897 | 0.07527 | 5 |
| 10 µM | Imidacloprid | 1 | ± | 0.1198 | 3 | 1.17751 | ± | 0.08629 | 0.26539 | 5 |
| 10 µM | Iprodione | 1 | ± | 0.07791 | 3 | 1.05483 | ± | 0.03381 | 0.50551 | 4 |
| 10 µM | Kelthane | 1 | ± | 0.21329 | 4 | 1.08744 | ± | 0.15685 | 0.75243 | 4 |
| 10 µM | Malathion | 1 | ± | 0.05808 | 3 | 0.80946 | ± | 0.09079 | 0.16702 | 4 |
| 10 µM | Mancozeb | 1 | ± | 0.06756 | 3 | 1.28397 | ± | 0.18586 | 0.26639 | 4 |
| 10 µM | Maneb | 1 | ± | 0.17309 | 4 | 0.69149 | ± | 0.1178 | 0.1634 | 6 |
| 10 µM | Mepiquat Chloride | 1 | ± | 0.12224 | 3 | 0.88203 | ± | 0.08977 | 0.45996 | 4 |
| 10 µM | Metalyxl | 1 | ± | 0.08271 | 3 | 0.96654 | ± | 0.11384 | 0.8445 | 5 |
| 10 µM | Metam Sodium | 1 | ± | 0.29137 | 3 | 1.59714 | ± | 0.08651 | 0.07349 | 4 |
| 10 µM | Methyl Bromide | 1 | ± | 0.07939 | 3 | 0.68226 | ± | 0.02165 | 0.00123 | 6 |
| 10 µM | Mevinphos | 1 | ± | 0.0726 | 3 | 0.88155 | ± | 0.04299 | 0.18005 | 5 |
| 10 µM | Napropamide | 1 | ± | 0.12796 | 4 | 1.41872 | ± | 0.0905 | 0.02471 | 6 |
| 10 µM | Norflurazon | 1 | ± | 0.06036 | 3 | 1.01852 | ± | 0.04307 | 0.80665 | 5 |
| 10 µM | Oxyfluorfen | 1 | ± | 0.05398 | 3 | 1.15783 | ± | 0.14397 | 0.41151 | 4 |
| 10 µM | Paraquat | 1 | ± | 0.03914 | 4 | 1.27593 | ± | 0.11678 | 0.06632 | 4 |
| 10 µM | Parathion Methyl | 1 | ± | 0.29167 | 3 | 1.26313 | ± | 0.16964 | 0.44292 | 4 |
| 10 µM | Pendimethalin | 1 | ± | 0.06066 | 3 | 1.20665 | ± | 0.13343 | 0.26792 | 4 |
| 10 µM | Phorate | 1 | ± | 0.15512 | 3 | 0.61918 | ± | 0.1055 | 0.08108 | 7 |
| 10 µM | Piperonyl Butoxide | 1 | ± | 0.04779 | 3 | 1.17469 | ± | 0.07731 | 0.14056 | 4 |
| 10 µM | Potassium Hydroxide | 1 | ± | 0.13255 | 3 | 0.74787 | ± | 0.04199 | 0.04953 | 6 |
| 10 µM | Prometryn | 1 | ± | 0.04626 | 3 | 1.26343 | ± | 0.02482 | 0.00289 | 4 |
| 1 µM | Rotenone | 1 | ± | 0.20067 | 4 | 0.44065 | ± | 0.03113 | 0.0331 | 4 |
| 10 µM | Sethoxydim | 1 | ± | 0.12809 | 3 | 1.0491 | ± | 0.05596 | 0.71277 | 4 |
| 10 µM | Sodium Arsenite | 1 | ± | 0.07238 | 4 | 1.36189 | ± | 0.12412 | 0.05159 | 5 |
| 10 µM | Sodium Cacodylate | 1 | ± | 0.13275 | 3 | 1.33045 | ± | 0.10012 | 0.09205 | 5 |
| 10 µM | Sodium Chlorate | 1 | ± | 0.04208 | 4 | 0.98064 | ± | 0.08892 | 0.86206 | 5 |
| 10 µM | Strychnine | 1 | ± | 0.19343 | 3 | 1.38805 | ± | 0.12732 | 0.13961 | 4 |
| 10 µM | Sulfur | 1 | ± | 0.06406 | 3 | 0.71613 | ± | 0.02849 | 0.0065 | 4 |
| 10 µM | Thiophanate Methyl | 1 | ± | 0.1231 | 3 | 0.97905 | ± | 0.09826 | 0.89799 | 4 |
| 10 µM | Triadimefon | 1 | ± | 0.1033 | 3 | 0.7852 | ± | 0.05606 | 0.10571 | 4 |
| 10 µM | Triflumizole | 1 | ± | 0.14608 | 3 | 1.55191 | ± | 0.09186 | 0.01979 | 4 |
| 10 µM | Trifluralin | 1 | ± | 0.10761 | 3 | 0.91044 | ± | 0.0577 | 0.46393 | 4 |
| 10 µM | Triforine | 1 | ± | 0.03308 | 3 | 1.02125 | ± | 0.06262 | 0.79887 | 4 |
| 10 µM | Vinclozolin | 1 | ± | 0.22631 | 3 | 1.6342 | ± | 0.25521 | 0.1353 | 4 |
| 10 µM | Zineb | 1 | ± | 0.04006 | 3 | 0.35255 | ± | 0.02058 | 2E-05 | 4 |
| 10 µM | Ziram | 1 | ± | 0.16916 | 5 | 2.34587 | ± | 0.52758 | 0.0525 | 6 |

**Supplement Table 3.**
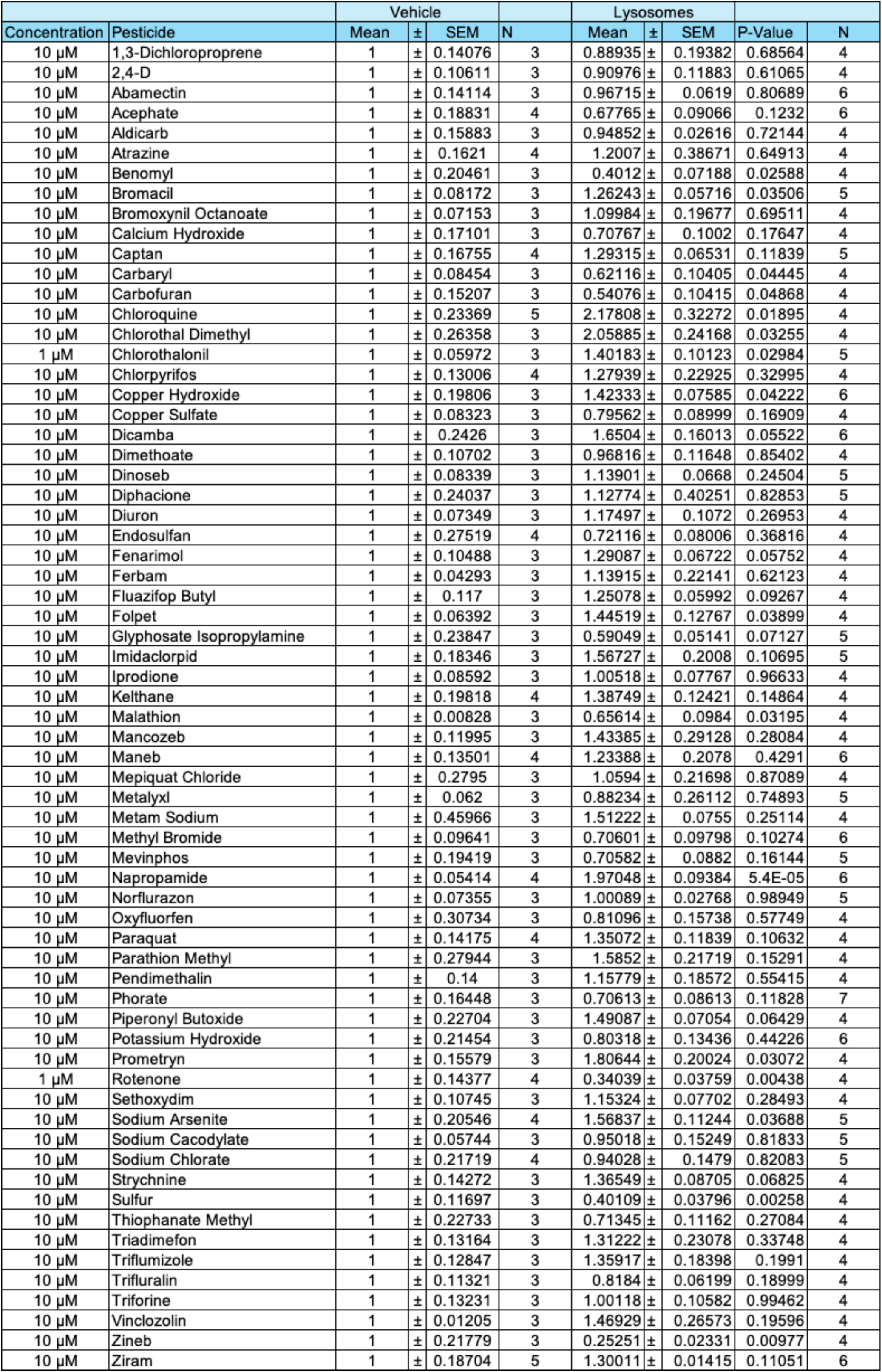
Results of lysosome foci differences in SK-N-MC autophagy assay. Pair-wise comparisons were conducted by Student’s T-test; raw foci counts for each pesticide was compared to a paired-vehicle condition. The pesticide is considered a hit at P<0.05.

**Supplement Figure 3.**
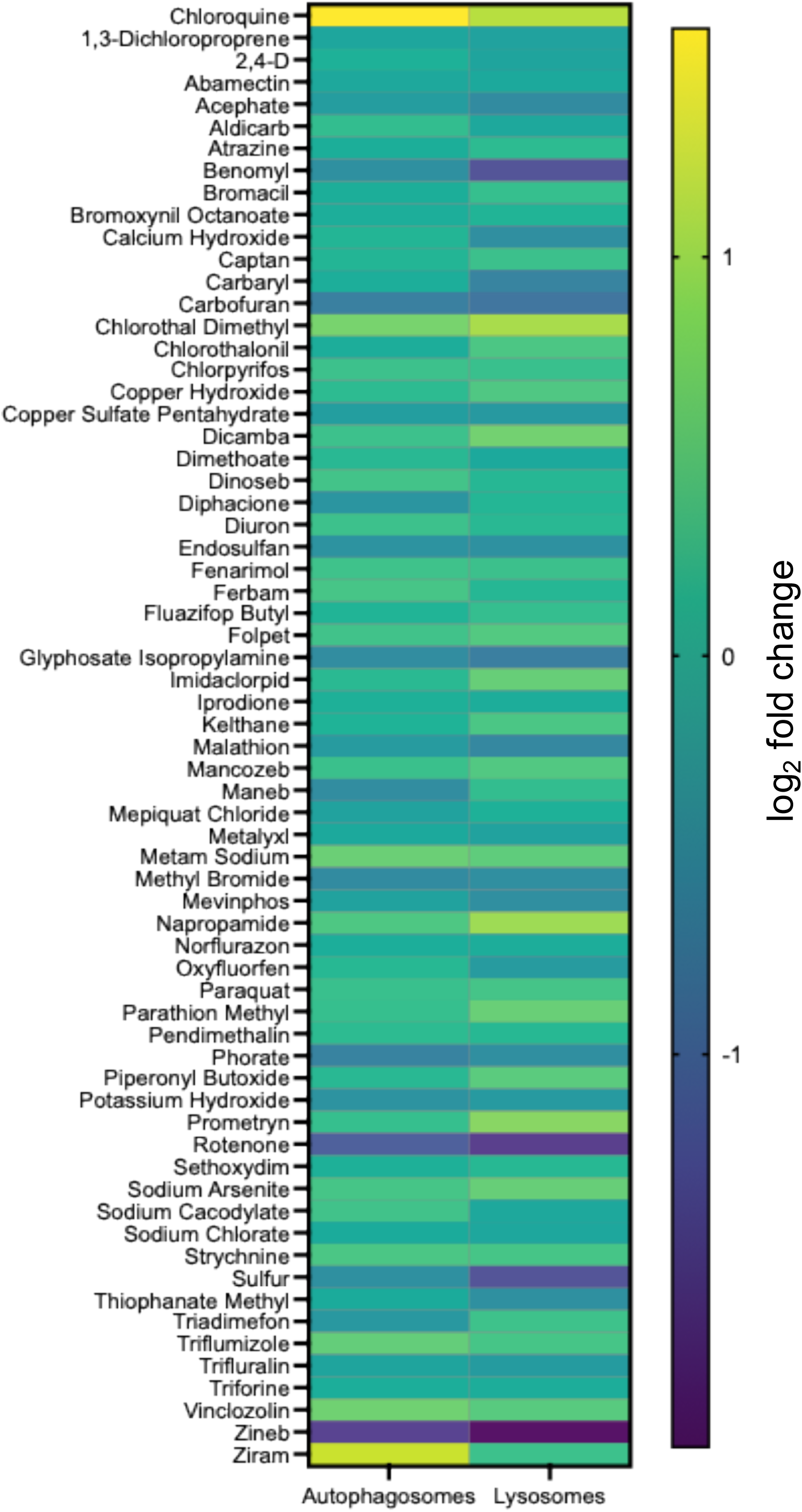
Effects of pesticides on autophagosome and lysosome counts in SK-N-MC cells. Data shown as log_2_ fold change relative to vehicle-treated cells following a 24- hour treatment with pesticide.

